# Activity of Sulfa Drugs and Their Combinations against Stationary Phase *B. burgdorferi* in vitro

**DOI:** 10.1101/112607

**Authors:** Jie Feng, Shuo Zhang, Wanliang Shi, Ying Zhang

**Affiliations:** Department of Molecular Microbiology and Immunology, Bloomberg School of Public Health, Johns Hopkins University, Baltimore, MD 21205, USA

**Keywords:** Borrelia burgdorferi, stationary phase cells, persisters, sulfa drugs, drug combination

## Abstract

Lyme disease is a most common vector borne disease in the US. Although the majority of Lyme patients can be cured with the standard 2-4 week antibiotic treatment, at least 10-20% of patients continue to suffer from prolonged post-treatment Lyme disease syndrome (PTLDS). While the cause for this is unclear, one possibility is that persisting organisms are not killed by current Lyme antibiotics. In our previous studies, we screened an FDA drug library and an NCI compound library on *B. burgdorferi* and found some drug hits including sulfa drugs as having good activity against *B. burgdorferi* stationary phase cells. In this study, we evaluated the relative activity of three commonly used sulfa drugs sulfamethoxazole (Smx), dapsone (Dps), sulfachlorpyridazine (Scp), and also trimethoprim (Tmp), and assessed their combinations with the commonly prescribed Lyme antibiotics for activities against *B. burgdorferi* stationary phase cells. Using the same molarity concentration, dapsone, sulfachlorpyridazine and trimethoprim showed very similar activity against stationary phase *B. burgdorferi* enriched in persisters, however, sulfamethoxazole was the least active drug among the three sulfa drugs tested. Interestingly, contrary to other bacterial systems, Tmp did not show synergy in drug combinations with the three sulfa drugs at their clinically relevant serum concentrations against *B. burgdorferi*. We found that sulfa drugs combined with other antibiotics were more active than their respective single drugs and that four-drug combinations were more active than three-drug combinations. Four drug combinations dapsone+minocycline+cefuroxime+azithromycin and dapsone+minocycline+cefuroxime+rifampin showed best activity against stationary phase *B. burgdorferi* in these sulfa drug combinations. However, these 4-sulfa drug containing combinations still had considerably less activity against *B. burgdorferi* stationary phase cells than the daptomycin+cefuroxime+doxycycline used as a positive control which completely eradicated *B. burgdorferi* stationary phase cells. Future studies are needed to evaluate and optimize the sulfa drug combinations *in vitro* and also in animal models.

## 1 Introduction

Lyme disease, which is caused by *Borrelia burgdorferi* sensu lato complex species, is the most common vector-borne disease in the United States with an estimated 300,000 cases a year [1]. The infection is transmitted to humans by tick vectors that feed upon rodents, reptiles, birds, deer, etc. [2]. In the early stage of Lyme disease, patients often have localized erythema migrans rash that expands as the bacteria disseminate from the cutaneous infection site via blood stream to other parts of the body. Late stage Lyme disease is a multi-system disorder which can cause arthritis and neurologic manifestations [1]. While the majority of Lyme disease patients can be cured if treated promptly with the standard 2-4 week doxycycline, amoxicillin, or cefuroxime therapy [3], at least 10-20% of Lyme patients have lingering symptoms such as fatigue, muscular and joint pain, and neurologic impairment even 6 months after the antibiotic treatment - a set of symptoms called Post-Treatment Lyme Disease Syndrome (PTLDS)[4] While the cause of PTLDS is unknown, several possibilities are likely to be involved, including autoimmune response [5], immune response to continued presence of antigenic debris [6], tissue damage as a result of Borrelia infection and inflammation, co-infections <u>[7]</u>, as well as persistent infection due to *B. burgdorferi* persisters that are not killed by the current antibiotics used to treat Lyme disease [8-10]. Various studies have found evidence of *B. burgdorferi* persistence in dogs [11], mice [8,9], monkeys [10], as well as humans [12] after antibiotic treatment, however, viable organisms are very hard to be cultured from the host after antibiotic treatment.

*B. burgdorferi* develops various forms of dormant persisters in stationary phase cultures which are tolerant to antibiotics used to treat Lyme disease [13-16]. These persister bacteria have an altered gene expression profile, which may underlie their drug tolerant phenotype [17]. In log phase cultures (3-5 day old), *B. burgdorferi* is primarily in motile spirochetal form which is highly susceptible to current Lyme antibiotics doxycycline and amoxicillin, however, in stationary phase cultures (7-15 day old), increased numbers of atypical forms such as round bodies and aggregated biofilm-like microcolonies develop [13,14]. These atypical forms have been shown to have increased tolerance to doxycycline and amoxicillin when compared to the growing spirochetal forms [13-16]. In addition, that the active hits from the round body persister screens [18] overlap with those from the screens on stationary phase cells [13] indicate the stationary phase culture or cells can be used as relevant persister model. Therefore, stationary phase cultures (7-15 day old) enriched in persisters have been used as a model for high-throughput drug screens against persisters [13,14,19].

We have recently identified a range of drugs with high activity against stationary phase cells enriched in persister through screens of FDA approved drug library and NCI compound libraries [13,19]. Besides daptomycin, clofazimine and cephalosporin antibiotics, we also found sulfa drugs as having good activity against *B. burgdorferi* stationary phase cells as well as round body forms [19]. In addition, a recent study found that sulfa drug dapsone had clinical benefit in the treatmnt of Lyme disease patients with persistent symptoms [20]. However, the relative activity of different sulfa drugs against *B. burgdorferi* stationary phase cells has not been evaluated in the same study under the same conditions. In this study, we compared the relative activity of three commonly used sulfa drugs sulfamethoxazole (Smx), dapsone (Dps), sulfachlorpyridazine (Scp) and trimethoprim (Tmp) (a drug that also inhibits folate pathway but is not considered a sulfa drug), and assessed their combinations with the commonly prescribed Lyme antibiotics and other antibiotics with activities against *B. burgdorferi* stationary phase cells as a persister model.

## 2 Results and Discussion

### 2.1. Comparison of the Relative Anti-Persister Activity of Commonly Used Sulfa Drugs Sulfamethoxazole, Dapsone, Sulfachlorpyridazine, and Trimethoprim for Their Activity against Stationary Phase B. burgdorferi Culture

To compare the activity of sulfamethoxazole, dapsone, sulfachlorpyridazine, and trimethoprim against *B. burgdorferi* stationary phase cells, we tested them on the same 7 day old *B. burgdorferi* stationary phase culture with the same molarity concentrations (10, 20, and 40 μM), using doxycycline and persister drug daptomycin as controls. Compared to the drug free control (residual viability 91%), the three sulfa drugs and trimethoprim (residual viability 88%-92%, Figure 1) showed little or no activity against stationary phase *B. burgdorferi* cultures at the low concentration (10 μM). Meanwhile, doxycycline control also showed no activity (residual viability 91%, Figure 1) against the stationary phase *B. burgdorferi.* At 10 μM concentration, we only found persister drug daptomycin showed good activity (residual viability 48%, Figure 1) against the stationary phase *B. burgdorferi* culture. To confirm results of the plate reader SYBR Green I/PI assay, we performed microscope counting SYBR Green I/PI assay on the antibiotic treated samples. The microscope counting results were in agreement with the plate reader results (Figure 1).

**Figure 1.**
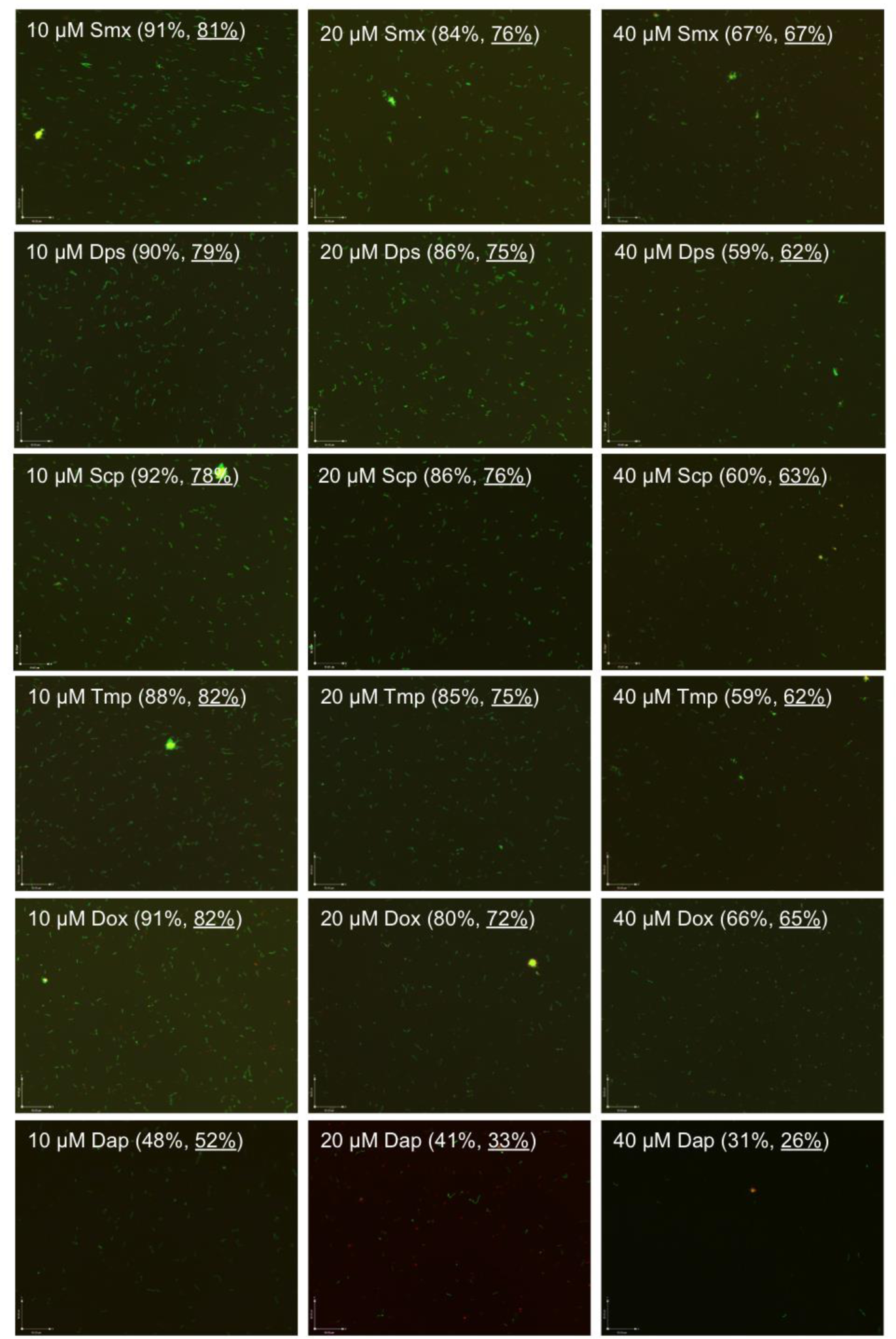
Comparison of anti-persister activity of sulfamethoxazole, dapsone, sulfachlorpyridazine and trimethoprim with doxycycline and daptomycin as controls. A 7-day old *B. burgdorferi* stationary phase culture containing aggregated microcolonies was incubated for 7 days with sulfamethoxazole (Smx), dapsone (Dps), sulfachlorpyridazine (Scp), trimethoprim (Tmp), doxycycline (Dox) and daptomycin (Dap) at the same drug concentrations of 10, 20 or 40 μM, respectively, followed by viability assessment using the SYBR Green I/PI assay. Representative images were taken using epifluorescence microscopy at 100× magnification. The calculated percentage and direct counting percentage (underlined) of residual viable cells is shown in brackets. The green cells stained by SYBR Green I dye indicate live cells while the red cells stained by PI dye indicate dead cells.

At the higher concentration (20 pM), the three sulfa drugs and trimethoprim had some activity (residual viability 86%-84%, Figure 1) against the *B. burgdorferi* stationary phase culture. We did not observe statistically significant difference in the activity of these four drugs. The three sulfa drugs and trimethoprim (residual viability 86%-84%) showed slightly less activity than doxycycline (residual viability 80%) but considerably less activity than daptomycin (residual viability 41%) at 20 μM concentration (Figure 1).

We found dose-dependent increase in killing activity of the three sulfa drugs and trimethoprim against the stationary phase *B. burgdorferi* resulting in 59%-67% residual viability (Figure 1) at the highest concentration (40 μM). Dapsone, sulfachlorpyridazine and trimethoprim showed very similar activity against stationary phase *B. burgdorferi* with the 59%, 60% and 59% residual viability respectively at 40 μM, however, sulfamethoxazole was the less active drug among the three sulfa drug and trimethoprim as shown by 67% residual viability after the drug treatment for 7 days (Figure 1). Meanwhile, we observed that dapsone, sulfachlorpyridazine and trimethoprim showed better activity than the doxycycline control (residual viability 66%) at the high concentration (40 μM). However, the three sulfa drugs and trimethoprim still could not eradicate stationary phase *B. burgdorferi* even at 40 μM. After 7 day drug treatment of sulfamethoxazole, dapsone, sulfachlorpyridazine and trimethoprim, we could still find many green (live) *B. burgdorferi* cells in aggregated microcolony form, round body form and spirochetal form under microscope revealed by the SYBR GreenI/PI viability assay (Figure 1). As shown in our previous studies [13,14,18], daptomycin showed impressive activity (residual viability 31%) against stationary phase *B. burgdorferi* at 40 μM as shown by mostly red (dead) cells and red microcolonies (Figure 1). Although dapsone, sulfachlorpyridazine, and trimethoprim showed better activity than doxycycline at 40 μM, their activity is relatively weak compared to daptomycin. We also noticed that sulfamethoxazole showed the weakest activity (residual viability 67%) among the sulfa drugs evaluated and is close to the activity of doxycycline (residual viability 66%) at the high concentration (40 μM).

### 2.2. Comparison of the Relative Anti-Persister Activity of Sulfamethoxazole, Dapsone, Sulfachlorpyridazine, and Trimethoprim in Drug Combinations at Respective Serum Drug Concentrations

Comparison with the same molar concentration of sulfamethoxazole, dapsone, sulfachlorpyridazine, and trimethoprim could reflect relative activity of these drugs; while testing the activity of these drugs and their drug combination at their respective serum concentration would provide clinically relevant information. To evaluate effective drug combinations that kill *B. burgdorferi* stationary phase culture at their serum concentrations, we tested sulfamethoxazole (15 μg/ml), dapsone (3 μg/ml), sulfachlorpyridazine (3 μg/ml), and trimethoprim (3 μg/ml) alone and their combinations with doxycycline (4 μg/ml), cefuroxime (5 μg/ml) and ciprofloxacin (3 μg/ml) on a 7 day old *B. burgdorferi* stationary phase culture. Using these clinically relevant concentrations, except sulfamethoxazole (residual viability 84%), sulfachlorpyridazine (residual viability 84%) and sulfamethoxazole+trimethoprim (residual viability 86%), all the other drugs or drug combinations showed killing activity compared to the drug free control (P<0.01, Figure 1) Sulfamethoxazole (residual viability 84%), dapsone (residual viability 83%), sulfachlorpyridazine (residual viability 84%), and trimethoprim (residual viability 84%) showed some killing activity compared to the drug free control (residual viability 95%) (Figure 1), but the differences between their activity were statistically insignificant (P>0.05).

Interestingly, trimethoprim (residual viability 84%) did not show synergy in the drug combinations with sulfamethoxazole (combination residual viability 86%), dapsone (combination residual viability 85%) and sulfachlorpyridazine (combination residual viability 86%). The results showed that the three sulfa drugs and trimethoprim combined with doxycycline (residual viability 85%), cefuroxime (residual viability 83%) and ciprofloxacin (residual viability 82%) were indeed more active (combination residual viability 74%-78%) than the single drugs (Table 1). However, we did not find significant difference of activity among these drug combinations. As shown in our previous studies [14,18], cefuroxime combined with doxycycline showed better activity (residual viability 78%) than them alone (Table 1). However, cefuroxime/doxycycline and cefuroxime/ciprofloxacin combined with sulfamethoxazole, dapsone, sulfachlorpyridazine or trimethoprim did not show higher activity (residual viability 74%-77%).

**Tabel 1.**
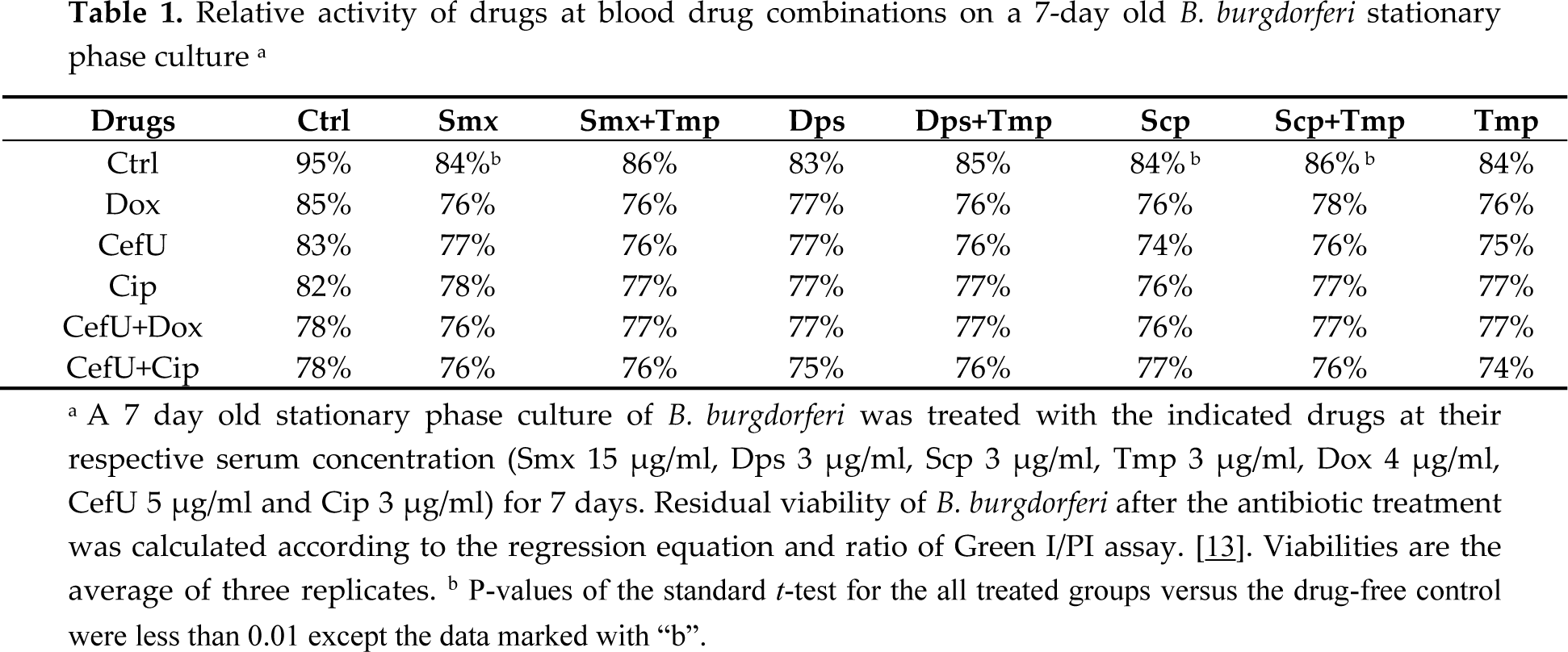
Relative activity of drugs at blood drug combinations on a 7-day old *B. burgdorferi* stationary phase culture ^a^

Our genomic analysis of *B. burgdorferi* B31 did not identify dihydropteroate synthase and dihydrofolate reductase, which respectively are the known targets of sulfa drugs and trimethoprim in other bacteria. Therefore, sulfa drugs and trimethoprim may work on *B. burgdorferi* B31 through some unknown pathways. This may explain why sulfa drugs and trimethoprim did not show synergistic effect against *B. burgdorferi* as would be expected with other bacteria but instead showed more activity when combined with doxycycline, cefuroxime and ciprofloxacin (Figure 2).

**Figure 2.**
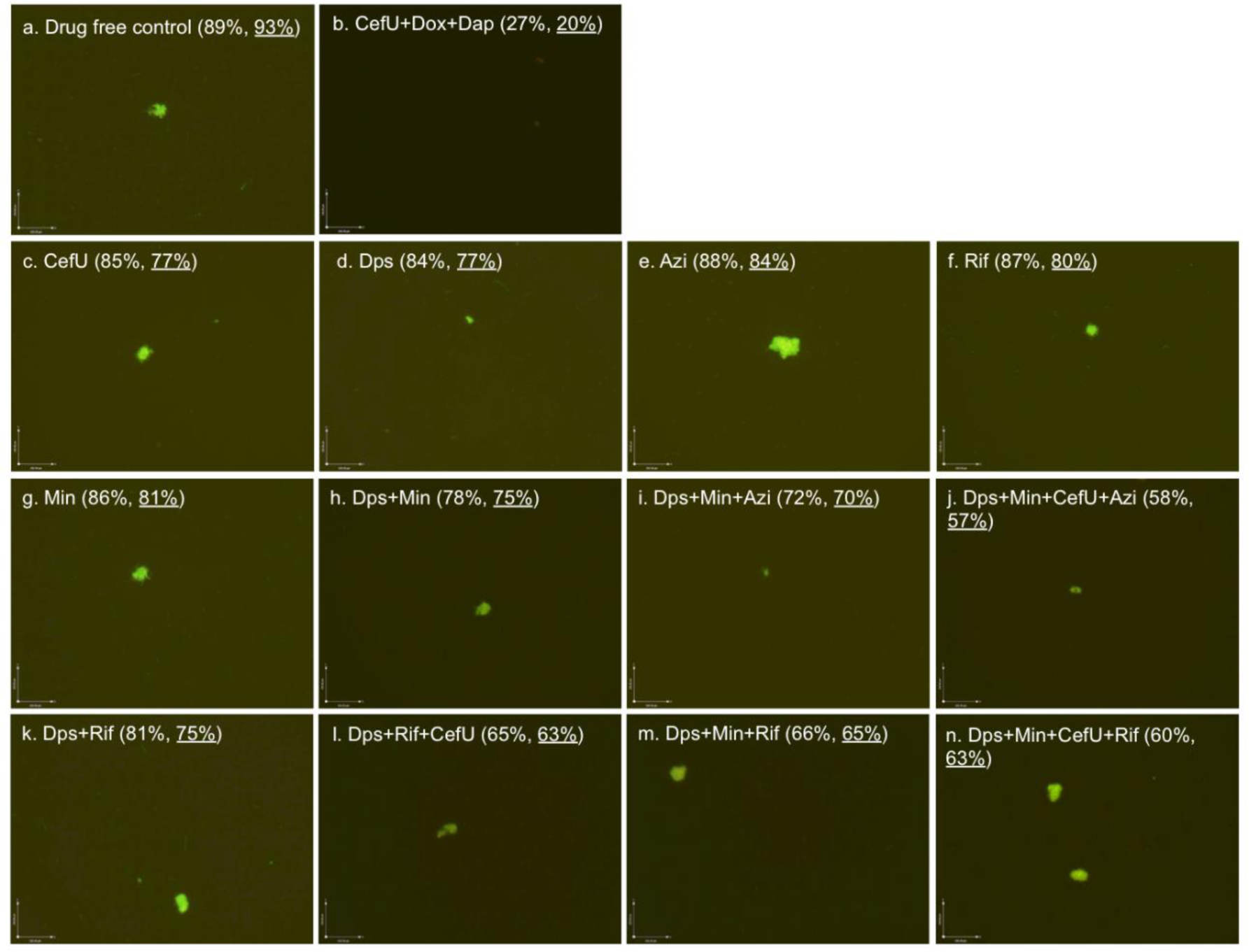
Effect of antibiotics alone or in combinations on stationary phase *B. burgdorferi* culture. A 7-day old *B. burgdorferi* stationary phase culture was incubated with the indicated drugs or drug combinations at a final concentration of 5 μg/mL for each antibiotic for 7 days followed by SYBR Green I/PI stain and epifluoresence microscopy (100× magnification). The calculated percentage and direct counting percentage (underlined) of residual viable cells is shown in brackets. Abbreviations: CefU, cefuroxime; Dps, dapsone; Azi, azithromycin; Rif, rifampin; Min, minocycline; Dox, doxycycline; Dap: daptomycin.

### 2.3. Effect of Dapsone Drug Combinations with Clinically Used Drugs on Stationary Phase B. burgdorferi Culture

Dapsone could improve chronic Lyme disease/PTLDS patients′ clinical symptoms in a recent study [20]. To identify more effective dapsone drug combinations that kill *B. burgdorferi* stationary phase cells, we tested some dapsone drug combinations with clinically used antibiotics (cefuroxime, azithromycin, rifampin and minocycline). The plate reader results were also confirmed with the microscope counting after SYBR Green I/PI viability staining (Figure 2). The results showed that some drug combinations were indeed much more effective than single drugs (Figure 2). Dapsone (residual viability 84%, Figure 2d) showed slightly better activity against the 7-day old stationary phase culture than the other three clinically used antibiotics azithromycin (residual viability 88%, Figure 2e), rifampin (residual viability 87%, Figure 2f) and minocycline (residual viability 86%, Figure 2g). Both two-drug combinations dapsone/minocycline and dapsone/rifampin showed better activity, with 78% and 81% residual viable (green) cells (Figure 2h and Figure 2k) remaining, respectively, in comparison to the single drugs (residual viability 84%-87%, Figure 2d, f and g). Interestingly, when azithromycin was added to the drug combination dapsone/rifampin, the anti-persister activity of these compounds was markedly increased as shown by 65% residual viable cells remaining (Figure 2j), compared to the dapsone/rifampin combination (residual viability 81%, Figure 2k). We also noted that rifampin/dapsone/minocycline showed better cooperative activity (residual viability 66%, Figure 2m) than the azithromycin/dapsone/minocycline combination (residual viability 72%, Figure 2i). Not surprisingly, four-drug combinations were more active (residual viability 58% and 60%, Figure 2j and n) than the three-drug combinations (residual viability 65%-72%, Figure 2i, l and n). In contrast to three-drug combination, azithromycin combined with dapsone/minocycline/cefuroxime was slightly more active (residual viability 58%, Figure 2j) than the rifampin four-drug combination (rifampin/dapsone/minocycline/cefuroxime, residual viability 60%, Figure 2n).

Sulfa drugs have recently been shown to have activity against *B. burgdorferi* pesisters[13,14] and more recently, dapsone has been shown to improve symptoms of patients with persistent Lyme disease [20]. Nevertheless, the relative activity of commonly used sulfa drugs such as sulfamethoxazole, dapsone and drug trimethroprim have not been compared under the same condition. In this study, we found that the different sulfa drugs dapsone, sulfachlorpyridazine, sulfamethoxazole and trimethoprim when used alone at their respective blood concentrations had similar but limited activity against *B. burgdorferi* stationary phase cells. However, at the same molar concentrations, trimethoprim had comparable activity as dapsone, both of which seem to be slightly more active than sulfachlorpyridazine and sulfamethoxazole (Figure 1).

Although sulfa drugs had some activity against *B. burgdorferi* stationary phase cells, their activities are enhanced when the sulfa drugs were combined with doxycycline, cefuroxime, ciprofloxacin, rifampin, or azithromycin, and the combination effects were more active than their respective single drugs. Among them, the oral four-drug combinations dapsone+minocycline+cefuroxime+azithromycin and dapsone+minocycline+cefuroxime+rifampin showed best activity (residual viable cells at 58% and 60%, respectively, Figure 2j, 2n) against stationary phase *B. burgdorferi*. However, the sulfa drug combinations containing even up to 4 drugs still had considerably less activity against *B. burgdorferi* stationary phase cells than the best drug combination daptomycin+cefuroxime+doxycycline (residual viable cells at 27%) used as a positive control (Figure 2b). As in our previous study [23], the daptomycin+cefuroxime+doxycycline drug combination could eradicate all *B. burgdorferi* cells as shown by all red (dead) cells under microscope Figure 2b).

In our previous studies [14,18,23], we used subculture to compare the samples with less than 30% viability after drug treatment to confirm whether the drugs eradicate the bacteria completely. However, in this study, most borrelia were still viable after sulfa drug or drug combination treatment with viable cells above 60%. In this case, recovery subculture would not find any difference between these samples, and therefore, we did not perform subculture tests in this study.

It is worth noting the present study is *in vitro* and as such the finding may have limitations. Levels of antibiotics in *in vitro* systems and the degree of protein binding in BSK medium are quite different from human serum, and it remains to be seen if the differences in relative activity of sulfa drugs and their combinations *in vitro* can be validated *in vivo* in animal models or in patients.

### 2.4 Effect of Sulfamethoxazole, Dapsone, Sulfachlorpyridazine, and Trimethoprim on Growing B. burgdorferi.

We also determined the minimum inhibitory concentration (MIC) of sulfamethoxazole, dapsone, sulfachlorpyridazine and trimethoprim on growing *B. burgdorferi* using the standard microdilution method. Our results showed that the three sulfa drugs and trimethoprim could partly inhibit the growth of *B. burgdorferi,* but cannot completely inhibit the growth even at 200 μg/mL (Figure 3). Sulfamethoxazole and trimethoprim showed obvious inhibition effect at lowest concentration (Figure 3c, g), and their combination nearly completely inhibited the growth of *B. burgdorferi* at 200 μg/mL(Figure 3l). However even the 200 μg/mL of these drugs still could not completely inhibit the growth of *B. burgdorferi* compared to the start culture (Figure 3d,f, h, l). This result is in disagreement with our previous test in which sulfamethoxazole showed a low MIC (less than 0.25 μg/mL) [13]. There are several possible reasons for the discrepant results. First, in this study we used animal passaged *B. burgdorferi* strain instead of previous unpassaged ATCC strain. Second, we could only selectively check some wells in the 96-well plate by counting chamber in the previous study, but in this study we checked every well directly in the 96-well plate with new BZ-X710 fluorescence microcopy. This greatly improved the experimental accuracy. In this study, we also found obvious growth inhibiting effect of sulfamethoxazole even at very low concentrations (0.4 μg/mL) (Figure 3c) compared to drug free control (Figure 3a). This inhibiting effect led to incorrect MIC determination because of the lack of day 0 start culture control in the previous study. Besides comparison to the drug free control (Figure 3b), sulfamethoxazole treated *B. burgdorferi* culture grew mainly in microcolony form instead of spirochetal form (Figure 3c,d) which could also have led to difficulties and inaccuracy of the counting chamber method used in the previous study. Meanwhile, consistent with our previous experiment [13], doxycycline and amoxicillin included as controls did not show growth of *B. burgdorferi* even at the lowest concentration of 0.2 μg/mL (Figure 3m-p).

**Figure 3.**
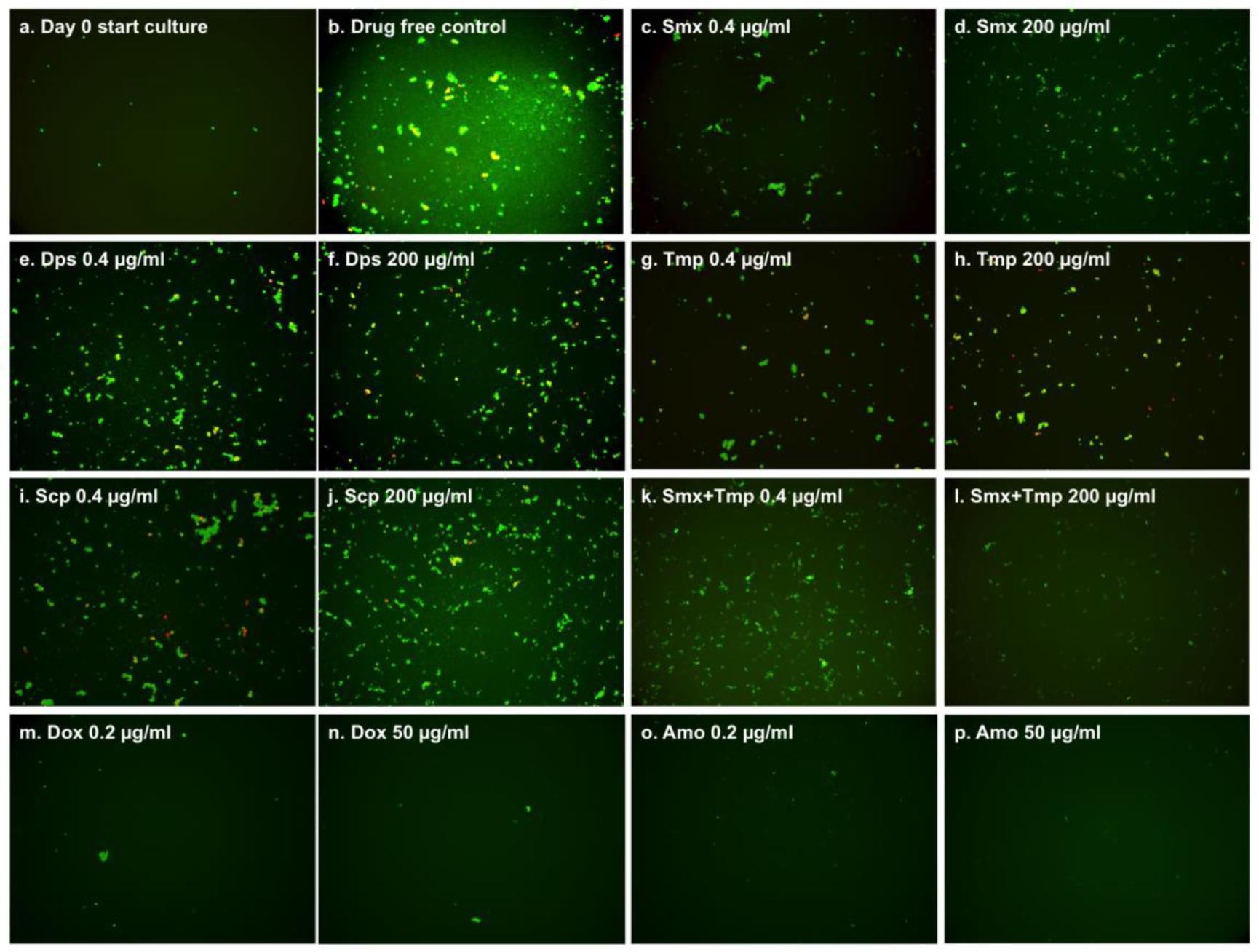
Effect of antibiotics on growing *B. burgdorferi* culture (5 day old). 10^4^ spirochetes in 90 μL fresh BSK-H medium as start culture (a) were incubated in 96-well microplate at 33°C and 5% CO_2_. The 10 μl 2-fold diluted antibiotics (200 μg/mL to 0.4 μg/mL) (c-l) and 10 μl DMSO as the drug free control (b) were added to the start culture. After 5-day incubation cell proliferation of every well was assessed using the SYBR Green I/PI assay and BZ-X710 All-in-One fluorescence microscope (KEYENCE, Inc.).

## 3. Experimental Section

### 3.1. Strain, Media and Culture Techniques

Low passaged *Borrelia burgdorferi* strain B31 5A19 was kindly provided by Dr. Monica Embers [16,21] The *B. burgdorferi* B31 strain was grown in BSK-H medium (HiMedia Laboratories Pvt. Ltd.) and supplemented with 6% rabbit serum (Sigma-Aldrich, St. Louis, MO, USA). All culture medium was filter-sterilized by 0.2 μm filter. Cultures were incubated in sterile 15 ml conical tubes (BD Biosciences, California, USA) in microaerophilic incubator (33°C, 5% CO2) without antibiotics. After incubation for 7-10 days, stationary-phase *B. burgdorferi* culture (about 10^7^ spirochetes/mL) was transferred into a 96 well plate for evaluation with the drugs or their combinations.

### 3.2. Drugs

The following drugs were obtained from Sigma-Aldrich, St. Louis, USA and dissolved in suitable solvents as suggested by the Clinical and Laboratory Standards Institute to make a 5 mg/mL stock solution: doxycycline (Dox), cefuroxime (CefU), ciprofloxacin (Cip), sulfamethoxazole (Smx), dapsone (Dps), sulfachlorpyridazine (Scp), trimethoprim (Tmp), azithromycin (Azi), rifampin (Rif), minocycline (Min) and daptomycin (Dap). The drug stock solutions were filter-sterilized using a 0.2 μm filter and stored at −20°C.

### 3.3. Microscopy

The *B. burgdorferi* cultures were examined using a Zeiss AxioImager M2 microscope with epifluorescence illumination. Pictures were taken using a SPOT slider camera. The SYBR Green I/PI viability assay was performed to assess cell viability using the ratio of green/red fluorescence to determine the live:dead cell ratio, respectively, as described previously [14]. This residual cell viability reading was confirmed by analyzing three representative images of the bacterial culture using epifluorescence microscopy. Image Pro-Plus software was used to quantitatively determine the fluorescence intensity.

### 3.4. Evaluation of Drugs and Drug Combinations for Their Activities against B. burgdorferi Stationary Phase Cultures

For assessing the activity of drugs and drug combinations against stationary phase *B. burgdorferi,* 5 μL aliquots of the drugs were added to 96 well plate containing 100 μL of the 7 day old stationary phase *B. burgdorferi* culture to obtain the desired drug concentration. Different drugs and drug combinations were evaluated at concentrations close to their Cmax values (maximum serum concentration). The plate was then sealed and was incubated at 33°C and 5% CO_2_ without shaking for 7 days when the residual viable cells remaining were calculated according to the regression equation and ratios of Green/Red fluorescence obtained by the SYBR Green I/PI viability assay, and then confirmed using epifluorescence microscopy as described [22]. Untreated groups were used as controls. Statistical analyses were performed using Student’s t-test. All experiments were run in triplicate.

### 3.5 Minimum inhibitory concentration (MIC) determination

The standard microdilution method was used to determine the MIC based on inhibition of visible growth of *B. burgdorferi* by microscopy. *B. burgdorferi* cells (1×10^4^) were inoculated into each well of a 96- well microplate containing 90 mL fresh BSK-H medium per well. Antibiotics were 2 fold diluted from 200 μg/mL to 0.4 μg/mL. Each diluted compound (10 μL) was added to the culture. All experiments were run in triplicate. The 96-well plate was sealed and placed in an incubator at 33°C for 5 days. Cell proliferation was assessed using the SYBR Green I/PI assay and BZ-X710 All-in-One fluorescence microscope (KEYENCE, Inc.) after the incubation.

## 4. Conclusions

In summary, dapsone, sulfachlorpyridazine and trimethoprim showed very similar activity against stationary phase *B. burgdorferi* at the same molarity concentration, and sulfamethoxazole was the least active drug among them. However, at blood concentrations, all the 4 drugs had similar activity. It is worth noting that trimethoprim did not show synergy in the drug combinations with the three sulfa drugs at their serum concentrations. However, sulfa drugs and trimethoprim when combined with other antibiotics such as doxycycline, ciprofloxacin and cefuroxime were more active than the respective single drugs. However, none of the sulfa drug combinations were as effective as daptomycin drug combination control since they were unable to completely eradicate *B. burgdorferi* stationary phase cells. Future studies are needed to optimize the drug combinations *in vitro* and to evaluate the sulfa drugs and their drug combinations *in vivo*.

## Acknowledgments

We acknowledge the support of our work by Steven & Alexandra Cohen Foundation, Global Lyme Alliance, Lyme Disease Association, and NatCapLyme. YZ was supported in part by NIH grants AI099512 and AI108535.

## Author Contributions

Ying Zhang conceived the experiments; Jie Feng, Wanliang Shi, Shuo Zhang, performed the experiments; Jie Feng and Ying Zhang analyzed the data; Jie Feng and Ying Zhang wrote the paper.

